# Allele-specific targeting of mutant TOR1A by the compact CRISPR/NmCas9 system in DYT1 dystonia with high fidelity

**DOI:** 10.1101/2024.04.22.590668

**Authors:** Junjiao Wu, Yu Tang

**Affiliations:** Department of Rheumatology and Immunology, Xiangya Hospital, Central South University, Changsha 410008, China; Provincial Clinical Research Center for Rheumatic and Immunologic Diseases, Xiangya Hospital, Central South University, Changsha 410008, China; National Clinical Research Center for Geriatric Disorders, Xiangya Hospital, Central South University, Changsha 410008, China; Aging Research Center, Department of Geriatrics, Xiangya Hospital, Central South University, Changsha 410008, China

**Keywords:** Dystonia, DYT1, TOR1A, Allele-specific, NmCas9

## Abstract

DYT1 is an autosomal dominant form of isolated dystonia, which is basically caused by an in-frame 3-bp GAG deletion in the *TOR1A* gene, leading to loss of a single glutamic acid residue (ΔE) at the C-terminus. TOR1A has been strongly implicated in various biological processes, such as protein quality control and regulation of ER stress. Many of these functions involve as protein multimers between TOR1A and its partners, whereas the ΔE mutant leads to destabilization of their binding, thereby reducing ATPase activation. Despite controversy over its functional model, the dominant-negative nature of TOR1A^ΔE^ has been demonstrated in a number of ways. Therefore, it is promising to develop an allele-specific intervention strategy that specifically silences the pathogenic TOR1A allele while preserving the wild-type allele to perform its normal function. In this study, we systematically evaluated the allele-specific targeting of TOR1A^ΔE^ using over 20 Cas endonucleases. We found that NmCas9, one of the compact Cas endonucleases yet with high-fidelity, selectively targeted the TOR1A^ΔE^ allele, with a 3-nt deletion located in the spacer region of sgRNAs. The discriminatory Nm-sgRNAs were verified both exogenously and endogenously that showed high specificity in disrupting the TOR1A^ΔE^ allele but not the wild-type one. Functionally, this strategy efficiently ameliorated the ubiquitin accumulation in DYT1 fibroblasts. Overall, our study demonstrates that the allele-specific targeting of mutant TOR1A with NmCas9 is a promising alternative approach for the treatment of DYT1.

## Introduction

DYT1 dystonia is an inherited form of isolated dystonia that manifests as involuntary movements or postural abnormalities due to sustained or intermittent muscle contractions ^1^. Genetically, this early-onset movement disorder is caused by a heterozygous 3-bp in-frame deletion in the last exon of the *TOR1A* gene (c.907_909delGAG), resulting in the loss of a single glutamic acid residue at position 302 or 303 from the TOR1A protein (TOR1A^ΔE^) ^2^. TOR1A is a ubiquitously expressed AAA+ protein localized in the contiguous lumen of the endoplasmic reticulum (ER)/nuclear envelope (NE) ^3^. To date, the biological role of TOR1A has yet fully delineated, however, it has been strongly implicated in a variety of biological functions, including control of protein quality as a molecular chaperone ^4–6^, ER stress related PERK-eIF2α signaling ^7,8^, regulation of linker of nucleoskeleton and cytoskeleton (LINC) complex and cell polarity ^9,10^, biogenesis of nuclear pores ^11,12^, membrane homeostasis and nucleocytoplasmic transport ^13,14^, synaptic vesicle cycling ^15,16^ and egress of large ribonucleoprotein granules through NE into the cytosol ^17,18^, among others.

Many of those functions may involve protein interactions in the ER. Accumulating evidence indicates that the TOR1A^ΔE^ mutation impairs TOR1A function in multiple ways. Essentially, AAA+ proteins function as multimers. TOR1A has been shown to interact with LAP1 (NE) and LULL1 (ER), respectively, to form a hexamer to trigger ATP hydrolysis and disassembly ^19,20^. This was also confirmed by crystallography ^21,22^. Specifically, the C-terminal region of TOR1A functions within the TOR1A/LULL1 and TOR1A/LAP1 hexamers, and also interacts with NESPRIN-3 ^9,23^, whereas the ΔE mutant causes destabilization of their binding, which reduces ATPase activation ^23^.

It is still controversy as to whether DYT1 dystonia is caused by loss-of-function (haploinsufficiency) or dominant-negative effects of the TOR1A^ΔE^ mutation. However, dominant-negative effects do participate in several pathological processes. For example, TOR1A^ΔE^ accumulates in NE, leading to the formation of NE-derived membranous inclusions known as spheroid bodies ^6,24,25^. Overexpression of TOR1A^ΔE^ leads to the recruitment of wild-type (WT) TOR1A to form spheroid bodies, which is suppressed by selectively silencing TOR1A^ΔE^ and thereby restoring its normal subcellular localization and functions ^26^. Similarly, TOR1A^ΔE^ inhibits the protein quality control mechanism, leading to aberrant accumulation of perinuclear ubiquitin ^27^. In addition, TOR1A facilitates the transit of replicating herpes simplex virus (HSV) capsids in the nucleus across the NE into the cytoplasm and out of the cell, a process that is interfered with by endogenous levels of TOR1A^ΔE^ ^28,29^. Truncation of the mutant allele by genome editing apparently enhances normal TOR1A activity and restores its function in promoting HSV replication in DYT1 fibroblasts ^30^. In further support of its dominant-negative effect, previous studies have also shown that selective silencing of mutant TOR1A mRNA using small interfering RNAs (siRNAs) or antisense oligonucleotides (ASOs) normalizes the DYT1 phenotype, and that TOR1A^ΔE^ acts in a dominant manner by inhibiting WT TOR1A activity ^26,31–35^.

Therefore, it is promising to develop an allele-specific targeting strategy that typically silences or ablates the pathogenic TOR1A allele, without compromising the normal function of the WT allele. Along with the development of CRISPR editing system, the allele-specific intervention turns from mRNA targeting to genomic targeting. Specifically, allele-specific CRISPR is able to discriminate between disease-causing alleles and WT alleles, depending on the incomplete complementarity of the sgRNA to the genomic sequence. The discriminatory sgRNAs, which provide the entry sites for Cas binding, contain critical elements including protospacer adjacent motif (PAM) sites and spacer sequences complementary to/targeting genetic variants. On this basis, allele-specific genome targeting is categorized into two working models: **‘*In PAM’*** and **‘*Near PAM’*** ^36^. Genetic variants or single nucleotide polymorphisms (SNPs) can create new PAM sites or eliminate PAM sites (**‘*In PAM’***), or they can be located within the seed region of spacer sequences (**‘*Near PAM’***), which renders their discrimination ability. The **‘*In PAM’*** model confers the most stringent allele-specific capabilities, because without a matching PAM, Cas binding to genomic DNA binding may be completely diminished. So far, allele-specific CRISPR has been exploited in treating various dominant diseases, such as retinitis pigmentosa, familial Alzheimer’s disease and dominant progressive hearing loss, as well as mutation-driven cancers ^36^.

In this study, we systematically evaluated allele-specific targeting of TOR1A^ΔE^ using more than 20 types of Cas endonucleases, based on both **‘*In PAM’*** and **‘*Near PAM’*** models. We found that NmCas9, one of the compact Cas endonucleases yet with high-fidelity ^37^, can selectively target TOR1A^ΔE^ allele, specifically due to a 3-nt deletion in the spacer region of Nm-sgRNAs. The discriminatory ability of Nm-sgRNAs was verified by exogenous and endogenous validation, and these Nm-sgRNAs showed a high degree of specificity in disrupting the TOR1A^ΔE^ allele but not the WT one. Notably, this allele-specific targeting efficiently produced functional outcomes that ameliorated the ubiquitin accumulations in DYT1 fibroblasts.

## Materials and Methods

### In silico analysis

We used the AsCRISPR website to select candidate sgRNA sequences for allele-specific editing (**Figure 1A; Supplementary Table 1**). Specifically, the following query sequence was input on the AsCRISPR website (http://www.genemed.tech/ascrispr) ^38^:

TGATGAAGACATTGTAAGCAGAGTGGCTGAG[GAG/-]ATGACATTTTTCCCCAAAGAGGAGAGAGTTT

**Figure 1.**
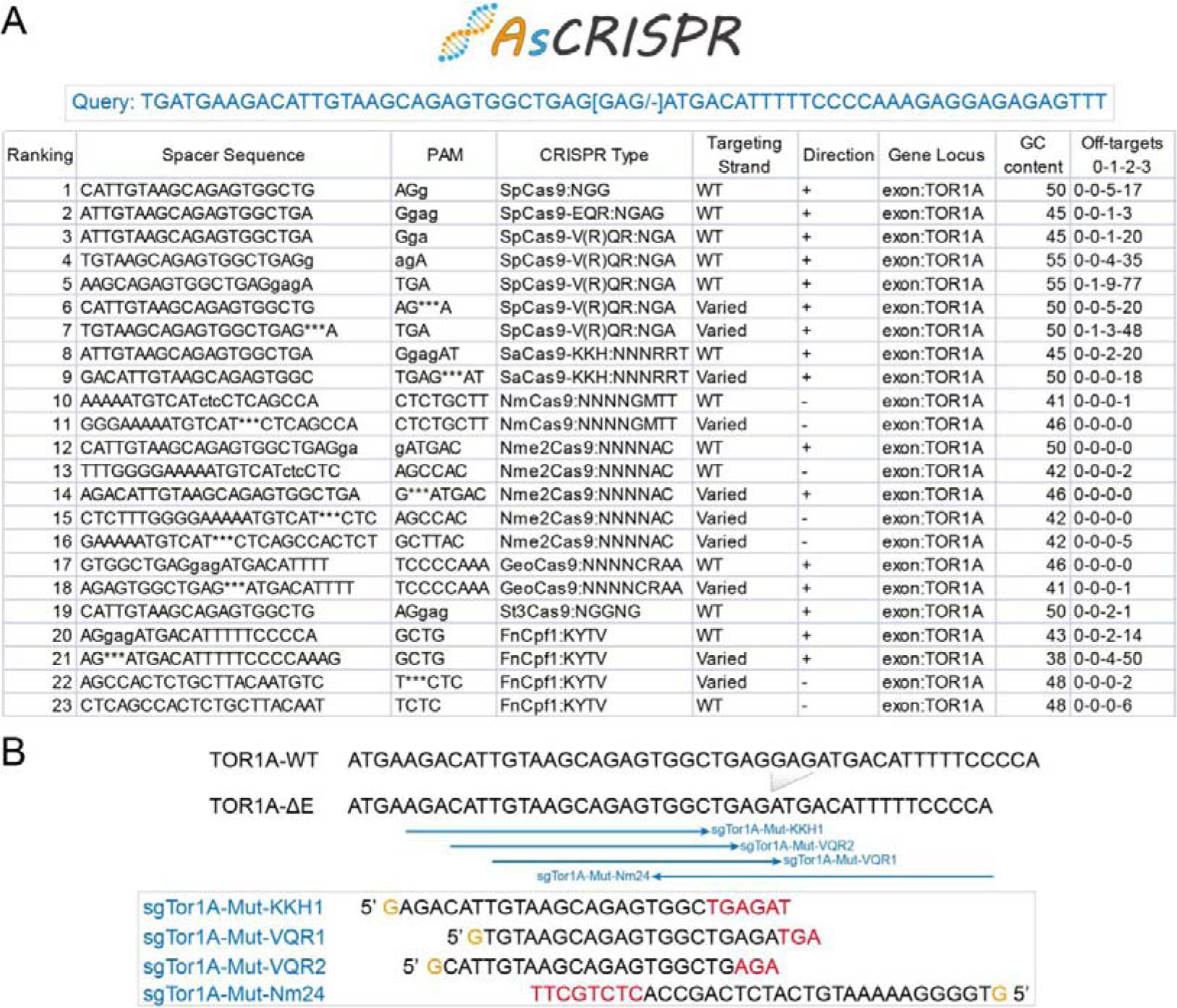
*In silico* analysis of allele-specific targeting. (A) Output of *in silico* analysis that comprehensively searched discriminatory sgRNAs and their paired Cas endonucleases, based on both ‘In PAM’ and ‘Near PAM’ models. (B) Four sgRNAs, working with SpCas9-VQR, SaCas9-KKH and NmCas9, respectively, were predicted to specifically target the mutant allele.

We then selected four major Cas endonucleases (Cas9, Cpf1, Cas12b and CasX), each of which contains several subtypes. A total of 24 Cas endonucleases, which recognize different PAM sites, were used for the analysis. The results output sgRNAs that may selectively target WT and mutant TOR1A and their potential off-targets. Candidate sgRNAs that may specifically target the mutant TOR1A were chosen (**Figure 1B**). Of those, sgRNAs of different lengths (18 to 24 nt) were designed for NmCas9 to evaluate the most efficient one (**Supplementary Table 1**).

### Constructions

To construct the reporter vector, pCAG-EGxxFP (addgene 50716) was used as the backbone. The sequences of TOR1A^WT^ and TOR1A^ΔE^ were amplified by PCR from the genomic DNA of a DYT1 fibroblast, and ligated into the BamHI (NEB) and EcoRI (NEB) digested sites of pCAG-EGxxFP. We constructed three CRISPR vectors, including lenti-U6-gRNA-NmCas9-HA-puro, lenti-U6-gRNA-SaCas9-KKH-puro and lentiCRISPR-SpCas9-VQR-Puro, for cloning sgRNAs.

Briefly, the U6-Nm tracrRNA fragment was amplified from pSimpleII-U6-sgRNA-NLS-NmCas9-HA-NLS (addgene 115694), and ligated into the KpnI/EcoRI linearized pXPR206-lenti-U6-gRNA-SaCas9-puro (addgene 96920), to generate lenti-U6-Nm gRNA scaffold-SaCas9-puro. The generated vector was further digested with AgeI/BamHI for use. Subsequently, the NmCas9-HA fragment was amplified and ligated to the digested vector by Gibson Assembly, which eventually produced the lenti-U6-gRNA-NmCas9-HA-puro vector.

The pXPR206-lenti-U6-gRNA-SaCas9-puro (addgene 96920) vector was linearized with AgeI/BamHI for use as the backbone. The SaCas9-FLAG fragment was then amplified from the SaCas9-KKH MSP1830 (addgene 70708) vector. The fragment and linearized vector were ligated by Gibson Assembly, to obtain lenti-U6-gRNA-SaCas9-KKH-puro. Similarly, the lentiCRISPR v2-Puro (addgene 52961) vector was digested with AgeI/BamHI for use. The FLAG-SpCas9-VQR fragment was amplified from the p459 SpCas9-VQR-Puro (addgene 101715) vector, and Gibson Assembly was performed to finally construct lentiCRISPR-SpCas9-VQR-Puro.

For cloning of sgRNAs, the three generated vectors served as backbones after BsmBI digestion. All designed sgRNA oligonucleotides were synthesized, denatured and annealed to form double-strand DNAs, which were respectively ligated to the linearized vectors using T4 ligase (Thermo). Sequences of sgRNAs and primers are listed in ***Supplementary Table 2***. All constructed plasmids were confirmed by Sanger sequencing before use.

### Cell culture

293T cells were routinely cultured with Dulbecco’s Modified Eagle Medium (DMEM; Gibco) supplemented with 10% fetal bovine serum (FBS, Biological Industries, Israel) and 1% each of penicillin / streptomycin (P/S) in a humidified incubator at 37°C with 5% CO_2_. Human primary fibroblasts were obtained from Coriell and NINDS-RUCDR, respectively (***Supplementary Table 3***). Three lines of DYT1 fibroblasts and six lines of normal fibroblasts were used in this study. All fibroblasts were cultured with DMEM containing 15% FBS and 1% P/S. The split ratios for passaging fibroblasts vary from 1:2 to 1:5 (***Supplementary Table 3***).

### The cleavage reporter assay

The constructed CRISPR vectors, as well as pCAG-EGxxFP vectors (containing WT and mutant TOR1A sequences), were combined separately and transfected into 293T cells using polyethylenimine (PEI, Polysciences). For transfection, 293T cells were passed and seeded at a density of 1.5 × 10^5^ cells per well in 12-well plates. The next day, 0.5 μg of each CRISPR plasmid and 0.5 μg of each pCAG-EGxxFP plasmid were added and mixed in OPTI-MEM (Gibco), followed by the addition of 3-fold volume of PEI. The mixture was allowed to stand at room temperature for 20 min, and then was slowly added to the culture medium. After 48 hrs of transfection, the medium was aspirated and the cells were washed once with PBS and replaced with fresh medium. EGFP fluorescence was observed and analyzed under a fluorescence microscope (Leica DMi8).

### Virus package and transduction

For virus package, 6 μg of each constructed vector, 4.5 μg of psPAX2 vector and 3 μg of PMD2.G vector were mixed in OPTI-MEM, and then 3-fold volume of PEI was added. The mixture was added into 293T cells in 10-cm dishes. After overnight, cells were rinsed twice with PBS and then replaced with fresh medium. After 48 and 72 hrs of transfection, the medium was harvested in centrifuge tubes and filtered through 0.45-μm filters (Millipore) to remove cell debris. The viruses were aliquoted and stored at −80°C until use. For transduction, fibroblasts were seeded into gelatin-coated T-25 culture flasks. The next day, viruses (MOI = 10) were added to the fibroblast culture medium with 6 μg/ml polybrene (Sigma). The cultures were replaced with fresh medium after 24 hrs. After three days, fibroblasts were passaged onto gelatin-coated coverslips for continual growth, till they were fixed with 4% paraformaldehyde (PFA) at room temperature for 20 min and ready for immunostaining.

### Generation of knockin cells

Four knockin 293T cell lines were generated: TOR1A^ΔE^-2A-BSD, TOR1A^WT^-2A-BSD, TOR1A^ΔE^-2A-GFP and TOR1A^WT^-2A-GFP. Briefly, sgRNA for KI (sgRNA-KI) was carefully designed by the CHOPCHOP webserver (http://chopchop.cbu.uib.no/) and cloned into lentiCRISPR v2-Puro. Notably, this sgTOR1A-KI did not have any predicted off-targets. The donor vectors were designed to contain 1000-bp genomic sequences flanking the cleavage site. Since the ΔE mutation is located at the 3’ end of the TOR1A coding region, we added blasticidin or GFP sequence after a 2A linker to form self-cleaving fusion proteins. This helps to screen knockin cells by antibiotic screening or cell sorting.

The flanking genomic sequences and 2A-BSD/GFP sequences were respectively amplified by PCR and together constructed into the SpeI/BamHI linearized pMD20T backbone by Gibson Assembly. To avoid re-cleavage, the PAM site (CGG) of the generated donor vectors were mutated into non-PAM (CGT) by site-directed mutagenesis. The sequence of sgTOR1A-KI and donor vectors are listed in ***Supplementary Table 4***.

To generate knockin cells, the lentiCRISPR v2-sgTOR1A-KI vector and different donor vectors were co-transfected into 293T cells using PEI. After 48 hrs, cells were treated with puromycin (1 μg/ml) for 24 hrs. Subsequently, to generate TOR1A^ΔE^-2A-BSD and TOR1A^WT^-2A-BSD knockin cells, the cell mixtures were passage with 0.25% Trypsin, and further diluted and seeded as 6000 single cells in a 10-cm dish. This allows for clonal cell line selection by manually picking single cell colonies for expansion. To generate TOR1A^ΔE^-2A-GFP and TOR1A^WT^-2A-GFP knockin cells, the cell mixtures were sorted by FACS (BD FACSCalibur) and further diluted as single cells for expansion. All picked colonies were characterized by junction PCR and Sanger sequencing. PCR primers are listed in ***Supplementary Table 4***.

### Cell proliferation and survival

For the cytotoxicity assay, cells were seeded at a density of 5 × 10^3^ cells/well on a 96-well plate for overnight, and were treated with blasticidin (20 μg/ml) on the next day. For the cell proliferation assay, cells were seeded at a density of 1 × 10^3^ cells/well on a 96-well plate. The cell viability/proliferation ability was assessed by MTT assay after 24, 48 and 72 hours, respectively. Briefly, 20 μL of MTT solution (5 mg/ml; Beyotime, China) was added into the wells and incubated at 37[ for 3 hrs. Then, the culture medium was carefully discarded, and the yielding formazan crystals were dissolved with 150 μl DMSO. Finally, ODs at 570 nm and 630 nm were measured and analyzed using a microplate reader (BioTek Cytation 5).

### Immunostaining

For immunostaining, fixed cells were permeabilized and blocked with the blocking solution containing 3% BSA and 0.3% Triton-X-100 at room temperature for 30 min. Cells were then incubated overnight at 4°C with primary antibodies, including HA (1:75; ABclonal AE008) and Ubiquitin (1:100; ABclonal A19686), respectively. The next day, cells were washed three times with PBST at a 10-min interval, and then incubated with the corresponding Alexa fluor-conjugated secondary antibodies (1:500; Invitrogen) at room temperature for 1 h. Again, cells were washed three times with PBST at a 10-min interval. Cells were counter-stained with DAPI for 5 min and then mounted with anti-fade Fluoromount-G (SouthernBiotech) for further observation under a fluorescence microscope (Leica DMi8).

### Western blot

For western blot, cells were harvested with RIPA buffer (Beyotime, China). Each sample was then added with 4 x loading buffer and β-ME and boiled at 95°C for 5 min. The proteins were separated by SDS-PAGE and transferred to a PVDF membrane (Millipore). The blots were blocked with 5% non-fat milk, and then incubated with diluted primary antibodies, including GFP (1:5000; Aves GFP-1020), Ubiquitin (1:800; ABclonal A19686), eIF2α (1:1000; Abclonal A0764), p-eIF2α (1:1000; Abclonal AP0692), PERK (1:1000; Abclonal A18196), p-PERK (1:800; Abclonal AP0886), ATF4 (1:1000; Abclonal A18687) and α-Tubulin (1:10000; Sigma T5168), at 4°C for overnight. On the next day, the blots were washed 3 times with TBST at a 15-min interval, and incubated with the HRP-conjugated secondary antibodies (1:5000; Servicebio, China) for 1 h at room temperature. After washing with TBST, western blots were detected using the ECL substrate and finally visualized by ChemiDoc XRS imager (Bio-Rad).

### Statistics analysis

GraphPad Prism software was utilized for statistical analysis and graphing. The Statistical analyses were performed using the Student’s t-test to determine the significance of differences between groups at the 95% confidence level. p-values less than 0.05 indicate statistical significance.

## Results

### In silico analysis of allele-specific targeting

To selectively target the TOR1A mutant allele, we first performed *in silico* analysis that comprehensively searched discriminatory sgRNAs, based on both **‘*In PAM’*** and **‘*Near PAM’*** models. Specifically, we evaluated four major types of Cas endonucleases including Cas9, Cpf1, Cas12b and CasX, as well as their subtypes, which recognize different PAM sequences. In this way, candidate sgRNAs and their paired Cas endonucleases were output (**Figure 1A**). In addition, a systematic evaluation of potential off-targets for each sgRNA was performed. It is noteworthy that no off-targets were predicted for Nm-sgRNAs, reminiscent of their high-fidelity. Four sgRNAs that were predicted to exclusively target the mutant allele were chosen for our study (**Figure 1B**). To take them into effect, SpCas9-VQR, SaCas9-KKH and NmCas9 were respectively employed.

### Verification of discriminatory sgRNAs

To verify the specificity of discriminatory sgRNAs, a cleavage reporter was constructed, based on the pCAG-EGxxFP vector, to involve a genomic fragment from either WT or mutant allele of TOR1A (**Figure 2A**). The discriminatory sgRNAs were also cloned into Cas9 containing vectors. Of note, we designed sgRNAs of different lengths (18 to 24 nt) for NmCas9 to evaluate the most efficient sgRNA (**Figure 2B**). Next, the constructed CRISPR vectors and reporter vectors were combined separately and transfected into 293T cells.

**Figure 2.**
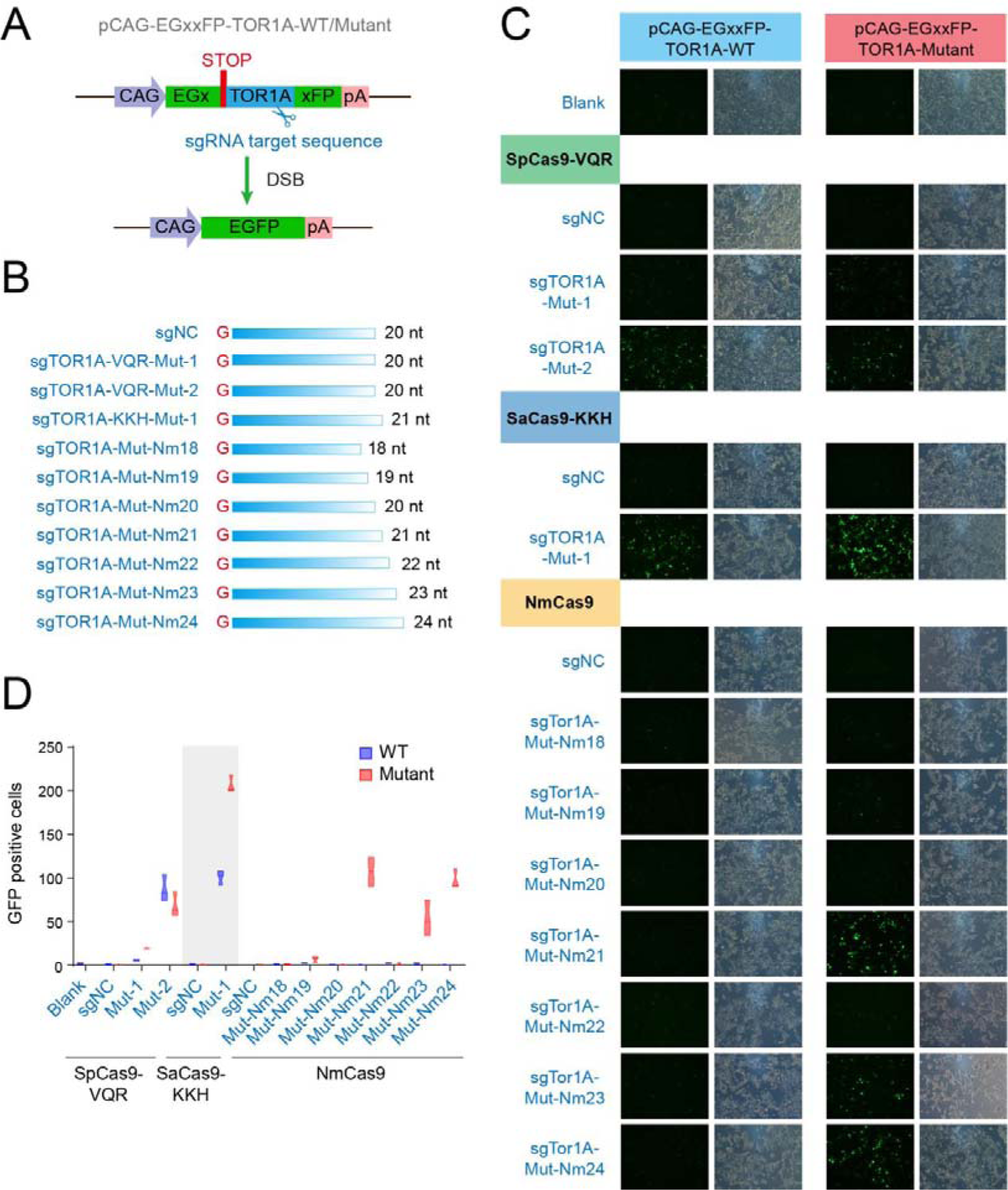
Verification of discriminatory sgRNAs. (A) The pCAG-EGxxFP reporter was constructed to include a TOR1A genomic fragment containing the sgRNA target sequence. The reporter contains 5’ and 3’ EGFP fragments with a 482-bp overlap. (B) discriminatory sgRNAs were cloned into Cas9 containing vectors. Different lengths (18 to 24 nt) of sgRNAs were designed for NmCas9 to assess the most efficient one. All sgRNAs were added with a G to their 5’ ends. (C, D) The CRISPR vectors, as well as reporter vectors, were combined separately and transfected into 293T cells. When the target sequence is cleaved by sgRNAs guided Cas9 endonuclease, the homology-dependent repair occurs, producing an intact GFP coding cassette. The cleavage effect can be assessed based on GFP production.

Specifically, if discriminatory sgRNAs are able to cleave the genomic DNA of TOR1A in the reporter vector, it triggers a homolog-dependent repair, including the homologous recombination of overlapped GFP sequences flanking the cleavage site, to produce a complete GFP coding cassette. Therefore, the cleavage efficacy can be assessed based on the production of GFP. The results showed that SpCas9-VQR/sgMut-VQR-2 was able to cleave both WT and mutant TOR1A with comparable efficiency, whereas sgMut-VQR-1 exhibited very limited capacity of DNA cleavage (**Figure 2C, D**). Interestingly, SaCas9-KKH/sgMut-KKH-1 were the most efficient in targeting the mutant allele, but they were also able to cleave the WT allele, albeit at half the efficiency (**Figure 2C, D**). While for NmCas9, sgMut-Nm21, sgMut-Nm23 and sgMut-Nm24 all selectively cleaved the mutant sequence to evoke GFP signals, but hardly functioned on the WT one (**Figure 2C, D**). Among them, the cleavage efficiency of sgMut-Nm23 was much lower. We thus choose sgMut-Nm21 and sgMut-Nm24 for further studies. Taken together, the results demonstrate that NmCas9/sgRNAs can be used for allele-specific targeting of TOR1A.

### Allele-specific targeting of endogenous TOR1A

To further evaluate the capabilities of NmCas9/sgRNAs in the allele-specific targeting of endogenous TOR1A mutant, we generated four knockin cell lines (TOR1A^ΔE^-2A-BSD, TOR1A^WT^-2A-BSD, TOR1A^ΔE^-2A-GFP and TOR1A^WT^-2A-GFP). Since the ΔE mutation resides on the 3’ terminus of TOR1A coding region, we added blasticidin or GFP sequence after a 2A linker to form self-cleaving fusion proteins (**Figure 3A**). Single cells were picked and expanded for characterization by junction PCR and western blot (**Figure 3B, C**). As confirmed by Sanger sequencing, we finally generated one homogenous TOR1A^ΔE^-2A-BSD strain, one heterogeneous TOR1A^WT^-2A-BSD strains, and one each strain of heterogeneous TOR1A^ΔE^-2A-GFP and TOR1A^WT^-2A-GFP (**Figure 3D**).

**Figure 3.**
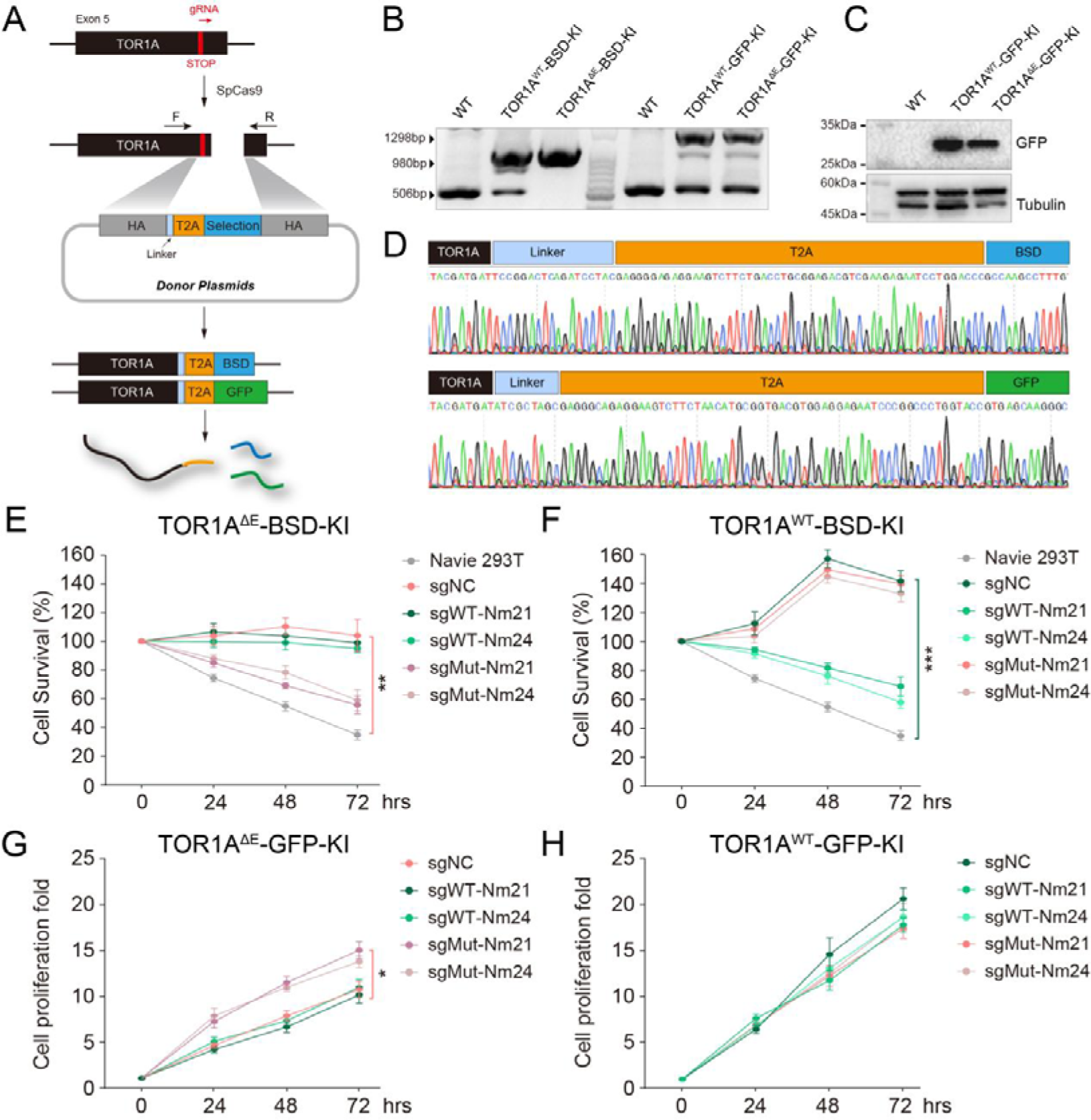
Allele-specific targeting of endogenous TOR1A. (A) Four knockin cell lines (TOR1A^ΔE^-2A-BSD, TOR1A^WT^-2A-BSD, TOR1A^ΔE^-2A-GFP and TOR1A^WT^-2A-GFP) were generated. Since the ΔE mutation is located at the 3’ end of TOR1A coding region, blasticidin or GFP sequence was added after a 2A linker to generate self-cleaving fusion proteins. (B, C) Single cells were picked and expanded for characterization by junction PCR and western blot. (D) Single cells were characterized by Sanger sequencing. (E, F) Assessment of resistance of TOR1A-2A-BSD cells to the blasticidin treatment, after allele-specific targeting. (G, H) Assessment of proliferative capacity of TOR1A-2A-GFP cells, after allele-specific targeting. *, p < 0.05; **, p < 0.01; ***, p < 0.001.

Later on, we tested the functional outcomes of allele-specific targeting in these knockin cells. Given that TOR1A-2A-BSD cells produce blasticidin, they are predicted to be more resistance to the blasticidin treatment. Indeed, both TOR1A^ΔE^-2A-BSD and TOR1A^WT^-2A-BSD survived well after the treatment. However, targeting by sgMut-Nm21/24 greatly diminished this resistance in TOR1A^ΔE^-2A-BSD cells (**Figure 3E**). As a control, TOR1A^WT^-2A-BSD cells were also vulnerable to the treatment after sgWT-Nm21/24 targeting (**Figure 3F**). It should be noted that although TOR1A^WT^-2A-BSD cells possessed only one copy of BSD, they showed to be more resistant to the blasticidin treatment, compared to TOR1A^ΔE^-2A-BSD cells that possessed two copies (**Figure 3F**). This may be due to the fact that the TOR1A^ΔE^ protein undergoes a stringent quality control by ER, leading to the hypo-expression of TOR1A^ΔE^ fused proteins, as also shown in **Figure 3C**.

We also found that allele-specific targeting of mutant TOR1A by sgMut-Nm21/24 effectively enhanced the proliferative capacity of TOR1A^ΔE^-2A-GFP cells (**Figure 3G**). However, targeting of the WT TOR1A by sgWT-Nm21/24 did not provide an additional impetus to the proliferation of TOR1A^WT^-2A-GFP cells, which were already growing much faster (**Figure 3H**). These results suggest that TOR1A^ΔE^ might execute a dominant-negative effect in regulating the cell cycle, which needs to be further validated.

TOR1A, as a molecular chaperone, plays an important role in protein quality and is actively involved in ER stress response. The quality control machinery was dampened by the mutant TOR1A, and caused abnormal accumulation of perinuclear ubiquitin ^27^. To investigate the effect of the mutant TOR1A in generated knockin cells, we then treated both heterogeneous TOR1A-2A-GFP cells with MG132, an inhibitor of the ubiquitin proteasome system (UPS), for analysis. There appeared to be no significant difference in PERK-eIF2α signaling associated with ER stress. However, under MG132 treatment, the TOR1A^ΔE^ mutant provoked stronger p-PERK/PERK and ATF4 signaling (**Supplementary Figure 1**). In addition, greatly increased ubiquitin accumulations were observed particularly in TOR1A^ΔE^-2A-GFP cells, reminiscent of an impaired quality control machinery (**Supplementary Figure 1**).

### Allele-specific targeting ameliorates the ubiquitin accumulation

To further test the functionality of discriminatory Nm-sgRNAs, we carried out the allele-specific targeting in human fibroblasts. We involved three DYT1 fibroblasts, which are all heterogeneous TOR1A^+/ΔE^, and six WT fibroblast controls (**Figure 4A**). Similar to those seen in knockin cells, we also detected increased expression of ubiquitin in DYT1 fibroblasts, as assessed by western blot (**Figure 4B, C**). Furthermore, by immunostaining, we could clearly observe the accumulation of perinuclear ubiquitin, which is also a pathological feature of DYT1 (**Figure 4D**).

**Figure 4.**
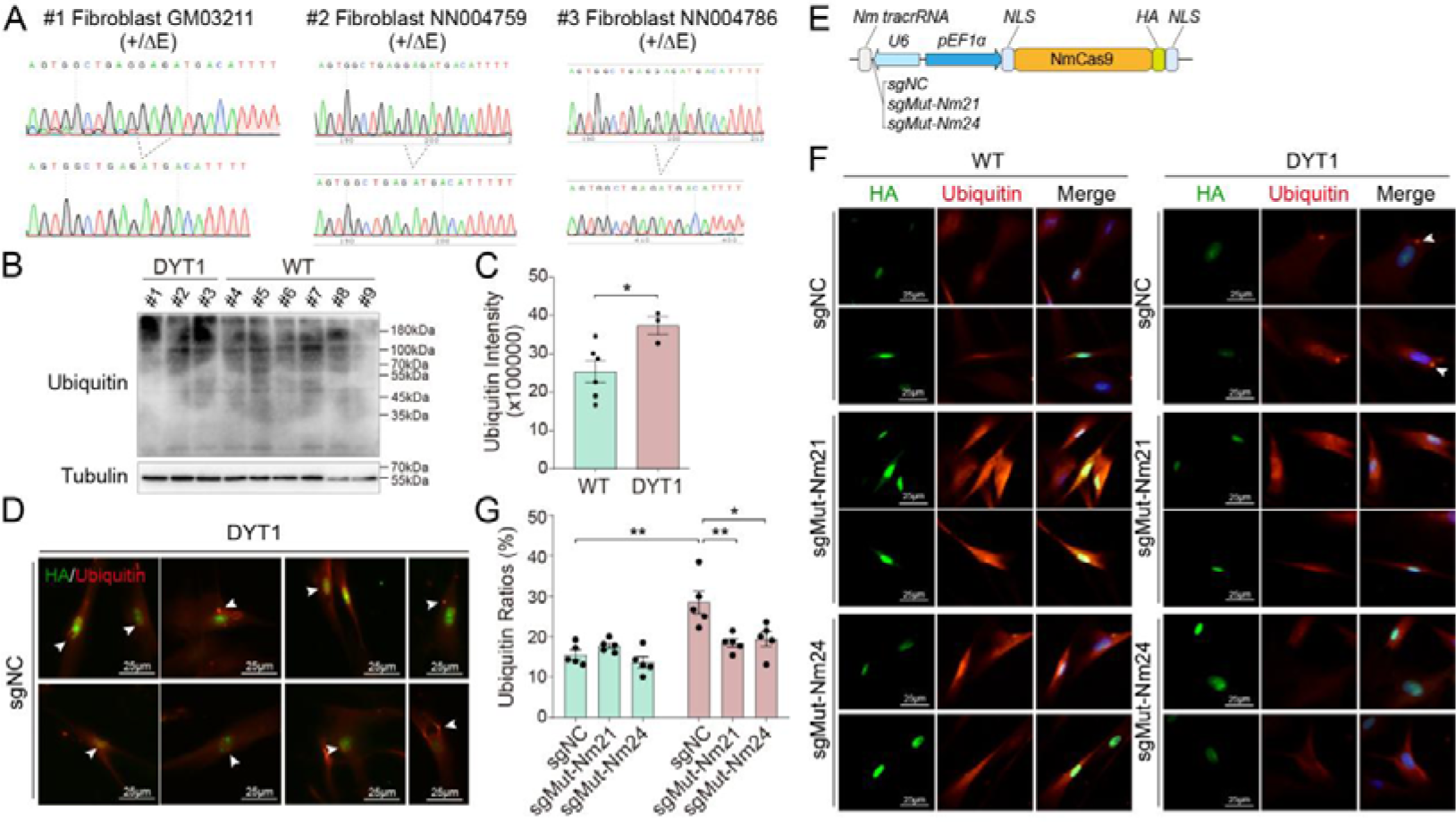
Allele-specific targeting ameliorates the ubiquitin accumulations in DYT1 fibroblasts. (A) Three DYT1 fibroblasts (heterogeneous TOR1A^+/ΔE^), and six WT fibroblast controls were involved. (B, C) Increased ubiquitin expression in DYT1 fibroblasts was assessed by western blot. (D) Prominent perinuclear ubiquitin accumulations were observed in DYT1 fibroblasts by immunostaining. Scale bars, 25 μm. (E) The NmCas9/sgNC, NmCas9/sgMut-Nm21 and NmCas9/sgMut-Nm24 vectors were packaged into lentiviruses for allele-specific targeting in fibroblasts, respectively. (F, G) The potential effects of discriminatory Nm-sgRNAs were examined by immunofluorescence staining. In DYT1 patient cells, the perinuclear ubiquitin accumulations were significantly reduced after targeting with sgMut-Nm21 and sgMut-Nm24. Scale bars, 25 μm. *, p < 0.05; **, p < 0.01.

We further packaged the NmCas9/sgNC, NmCas9/sgMut-Nm21 and NmCas9/sgMut-Nm24 vectors into lentiviruses, respectively, and transduced them into these fibroblasts (**Figure 4E**). The discriminatory effects of Nm-sgRNAs after allele-specific targeting were examined by immunofluorescence staining. The results first showed that this method was quite effective, since the transfection efficiency reached about 80% in both WT and DYT1 fibroblasts, as evidenced by the HA expression of CRISPR vectors and the GFP expression of control vectors (**Supplementary Figure 2**). In DYT1 patient cells, perinuclear ubiquitin accumulations were prominent in the sgNC group (**Figure 4D, F**), which were significantly reduced after targeting with sgMut-Nm21 and sgMut-Nm24 (**Figure 4F, G**). Overall, the results demonstrate that discriminatory NmCas9/sgMut-Nm21/24 holds the potential to attenuate the pathological features of DYT1 dystonia.

## Discussion

In this study, we systematically explored sgRNAs that effectively mediate their cleavage of genomic DNAs in an allele-specific manner. Specifically, the combination of NmCas9 with either sgMut-Nm21 or sgMut-Nm24 was validated to be highly specific in targeting the TOR1A^ΔE^ allele and produced beneficial effects in mitigating DYT1 pathology. Of note, one previous study has successfully disrupted the TOR1A mutant allele by SpCas9-VRQR, in which its sgRNA (H3 gRNA) performed pretty well (∼80% indel for the mutant allele, ∼20% indel for the WT allele), which was evaluated by deep sequencing ^30^. Interestingly, the sequence of H2 gRNA in that study (listed in ***Supplementary Table 2***) is identical to that of our designed sgMut-VQR-2, which we have shown to have limited discriminatory ability. This is in line with their results that H2 gRNA was shown to be 6-fold less efficient than the H3 gRNA ^30^. The only one 5’ nucleotide difference between H2 (GN_20_) and H3 (GN_19_) gRNA produced distinct cleavage effects, demonstrating that a candidate discriminating sgRNA should be carefully designed and thoroughly examined to achieve maximum benefits.

We assumed that another strength of this study is that NmCas9 is a quite compact Cas endonuclease (1082 aa), yet with higher fidelity and diminished off-targets ^37^, compared to the traditionally used SpCas9 (1368 aa). This renders NmCas9 to be more competent for packaging into adeno-associated viruses (AAVs), thereby boosting its translational capability of carrying out allele-specific targeting in DYT1 patients.

While NmCas9/sgMut-Nm21/24 may possess lower cleavage efficiencies than SaCas9-KKH, and probably SpCas9, other avenues may be used to increase the specificity of SaCas9 and SpCas9 when applied for allele-specific purposes. For example, a second mismatch might be added to the spacer region of discriminatory sgRNAs, which would result in two mismatched sgRNAs with undesired allele ^39^. However, given the compact size of NmCas9, increasing the cleavage capability of NmCas9 shall produce more promising advantages. To this end, more types of Cas endonucleases with expanded PAM sites are urgently required to be developed.

Alternatively, other CRISPR tools, such as base editors ^40–42^ or Cas13d ^43,44^, are also expected to achieve allele-specific effects. Notably, Cas13d has the unique strength that it specifically cleavage RNAs, rather than DNAs, and will not cause irreversible DNA changes to the cells ^43^. Rationally, it resembles RNA inference (RNAi), which has been widely used and developed for therapeutic applications ^45^. Importantly, allele-specific RNAi of TOR1A^ΔE^ has been investigated in several studies ^26,31–35^, rendering the Cas13d-mediated allele-specific targeting more practical.

As we applied the allele-specific endeavors in diseases, largely due to the fact that the genetic mutations act in a dominant-negative manner. The intervention outcomes are thus valuable for echoing their known effects, while also revealing the undiscovered mechanisms. By examining the discriminatory ability of NmCas9/sgMut-Nm21/24, several DYT1 pathological features were ameliorated; yet a possible dominant-negative role of TOR1A in regulating the cell cycle is worth noting. In addition, since DYT1 is recognized as a neural network disease ^46,47^, allele-specific targeting could also be applied to restore neuron-specific hypo-functions in DYT1-iPSCs and their derived neurons ^13,48–51^.

## Authors Contributions

YT conceived and designed the experiments. YT and JJW prepared the draft manuscript and figures. All authors have reviewed and approved the final version of the manuscript.

## Acknowledgements

We thank the members of the Tang laboratory for insightful discussions.

## References

1. Balint B, Mencacci NE, Valente EM, et al. Dystonia. Nat Rev Dis Primers. 2018;4(1):25.

2. Ozelius LJ, Hewett JW, Page CE, et al. The early-onset torsion dystonia gene (DYT1) encodes an ATP-binding protein. Nat Genet. 1997;17(1):40–48.

3. Goodchild RE, Kim CE, Dauer WT. Loss of the dystonia-associated protein torsinA selectively disrupts the neuronal nuclear envelope. Neuron. 2005;48(6):923–932.

4. Nery FC, Armata IA, Farley JE, et al. TorsinA participates in endoplasmic reticulum-associated degradation. Nat Commun. 2011;2:393.

5. Chen P, Burdette AJ, Porter JC, et al. The early-onset torsion dystonia-associated protein, torsinA, is a homeostatic regulator of endoplasmic reticulum stress response. Hum Mol Genet. 2010;19(18):3502–3515.

6. Goodchild RE, Dauer WT. Mislocalization to the nuclear envelope: an effect of the dystonia-causing torsinA mutation. Proc Natl Acad Sci U S A. 2004;101(3):847–852.

7. Rittiner JE, Caffall ZF, Hernandez-Martinez R, et al. Functional Genomic Analyses of Mendelian and Sporadic Disease Identify Impaired eIF2alpha Signaling as a Generalizable Mechanism for Dystonia. Neuron. 2016;92(6):1238–1251.

8. Beauvais G, Bode NM, Watson JL, et al. Disruption of Protein Processing in the Endoplasmic Reticulum of DYT1 Knock-in Mice Implicates Novel Pathways in Dystonia Pathogenesis. J Neurosci. 2016;36(40):10245–10256.

9. Nery FC, Zeng J, Niland BP, et al. TorsinA binds the KASH domain of nesprins and participates in linkage between nuclear envelope and cytoskeleton. J Cell Sci. 2008;121(Pt 20):3476–3486.

10. Starr DA, Rose LS. TorsinA regulates the LINC to moving nuclei. J Cell Biol. 2017;216(3):543–545.

11. VanGompel MJ, Nguyen KC, Hall DH, Dauer WT, Rose LS. A novel function for the Caenorhabditis elegans torsin OOC-5 in nucleoporin localization and nuclear import. Mol Biol Cell. 2015;26(9):1752–1763.

12. Rampello AJ, Laudermilch E, Vishnoi N, et al. Torsin ATPase deficiency leads to defects in nuclear pore biogenesis and sequestration of MLF2. J Cell Biol. 2020;219(6).

13. Ding B, Tang Y, Ma S, et al. Disease Modeling with Human Neurons Reveals LMNB1 Dysregulation Underlying DYT1 Dystonia. J Neurosci. 2021;41(9):2024–2038.

14. Yu-Taeger L, Gaiser V, Lotzer L, et al. Dynamic nuclear envelope phenotype in rats overexpressing mutated human torsinA protein. Biol Open. 2018;7(7).

15. Granata A, Watson R, Collinson LM, Schiavo G, Warner TT. The dystonia-associated protein torsinA modulates synaptic vesicle recycling. J Biol Chem. 2008;283(12):7568–7579.

16. Kakazu Y, Koh JY, Ho KW, Gonzalez-Alegre P, Harata NC. Synaptic vesicle recycling is enhanced by torsinA that harbors the DYT1 dystonia mutation. Synapse. 2012;66(5):453–464.

17. Speese SD, Ashley J, Jokhi V, et al. Nuclear envelope budding enables large ribonucleoprotein particle export during synaptic Wnt signaling. Cell. 2012;149(4):832–846.

18. Jokhi V, Ashley J, Nunnari J, et al. Torsin mediates primary envelopment of large ribonucleoprotein granules at the nuclear envelope. Cell Rep. 2013;3(4):988–995.

19. Chase AR, Laudermilch E, Wang J, Shigematsu H, Yokoyama T, Schlieker C. Dynamic functional assembly of the Torsin AAA+ ATPase and its modulation by LAP1. Mol Biol Cell. 2017;28(21):2765–2772.

20. Goodchild RE, Dauer WT. The AAA+ protein torsinA interacts with a conserved domain present in LAP1 and a novel ER protein. J Cell Biol. 2005;168(6):855–862.

21. Demircioglu FE, Zheng W, McQuown AJ, et al. The AAA + ATPase TorsinA polymerizes into hollow helical tubes with 8.5 subunits per turn. Nat Commun. 2019;10(1):3262.

22. Demircioglu FE, Sosa BA, Ingram J, Ploegh HL, Schwartz TU. Structures of TorsinA and its disease-mutant complexed with an activator reveal the molecular basis for primary dystonia. Elife. 2016;5.

23. Zhao C, Brown RS, Chase AR, Eisele MR, Schlieker C. Regulation of Torsin ATPases by LAP1 and LULL1. Proc Natl Acad Sci U S A. 2013;110(17):E1545–1554.

24. Naismith TV, Heuser JE, Breakefield XO, Hanson PI. TorsinA in the nuclear envelope. Proc Natl Acad Sci U S A. 2004;101(20):7612–7617.

25. Gonzalez-Alegre P, Paulson HL. Aberrant cellular behavior of mutant torsinA implicates nuclear envelope dysfunction in DYT1 dystonia. J Neurosci. 2004;24(11):2593–2601.

26. Gonzalez-Alegre P, Bode N, Davidson BL, Paulson HL. Silencing primary dystonia: lentiviral-mediated RNA interference therapy for DYT1 dystonia. J Neurosci. 2005;25(45):10502–10509.

27. Liang CC, Tanabe LM, Jou S, Chi F, Dauer WT. TorsinA hypofunction causes abnormal twisting movements and sensorimotor circuit neurodegeneration. J Clin Invest. 2014;124(7):3080–3092.

28. Gyorgy B, Cruz L, Yellen D, et al. Mutant torsinA in the heterozygous DYT1 state compromises HSV propagation in infected neurons and fibroblasts. Sci Rep. 2018;8(1):2324.

29. Turner EM, Brown RS, Laudermilch E, Tsai PL, Schlieker C. The Torsin Activator LULL1 Is Required for Efficient Growth of Herpes Simplex Virus 1. J Virol. 2015;89(16):8444–8452.

30. Cruz L, Gyorgy B, Cheah PS, et al. Mutant Allele-Specific CRISPR Disruption in DYT1 Dystonia Fibroblasts Restores Cell Function. Mol Ther Nucleic Acids. 2020;21:1–12.

31. Gonzalez-Alegre P, Miller VM, Davidson BL, Paulson HL. Toward therapy for DYT1 dystonia: allele-specific silencing of mutant TorsinA. Ann Neurol. 2003;53(6):781–787.

32. Kock N, Allchorne AJ, Sena-Esteves M, Woolf CJ, Breakefield XO. RNAi blocks DYT1 mutant torsinA inclusions in neurons. Neurosci Lett. 2006;395(3):201–205.

33. Hewett JW, Nery FC, Niland B, et al. siRNA knock-down of mutant torsinA restores processing through secretory pathway in DYT1 dystonia cells. Hum Mol Genet. 2008;17(10):1436–1445.

34. Martin JN, Wolken N, Brown T, Dauer WT, Ehrlich ME, Gonzalez-Alegre P. Lethal toxicity caused by expression of shRNA in the mouse striatum: implications for therapeutic design. Gene Ther. 2011;18(7):666–673.

35. Beauvais G, Watson JL, Aguirre JA, Tecedor L, Ehrlich ME, Gonzalez-Alegre P. Efficient RNA interference-based knockdown of mutant torsinA reveals reversibility of PERK-eIF2alpha pathway dysregulation in DYT1 transgenic rats in vivo. Brain Res. 2019;1706:24–31.

36. Wu J, Tang B, Tang Y. Allele-specific genome targeting in the development of precision medicine. Theranostics. 2020;10(7):3118–3137.

37. Amrani N, Gao XD, Liu P, et al. NmeCas9 is an intrinsically high-fidelity genome-editing platform. Genome Biol. 2018;19(1):214.

38. Zhao G, Li J, Tang Y. AsCRISPR: A Web Server for Allele-Specific Single Guide RNA Design in Precision Medicine. The CRISPR Journal. 2020;3(6):512–522.

39. Chen X, Zhao D, Hou X, et al. Genomic allele-specific base editing with imperfect gRNA. J Genet Genomics. 2023.

40. Komor AC, Kim YB, Packer MS, Zuris JA, Liu DR. Programmable editing of a target base in genomic DNA without double-stranded DNA cleavage. Nature. 2016;533(7603):420–424.

41. Gaudelli NM, Komor AC, Rees HA, et al. Programmable base editing of A*T to G*C in genomic DNA without DNA cleavage. Nature. 2017;551(7681):464–471.

42. Kim YB, Komor AC, Levy JM, Packer MS, Zhao KT, Liu DR. Increasing the genome-targeting scope and precision of base editing with engineered Cas9-cytidine deaminase fusions. Nat Biotechnol. 2017;35(4):371–376.

43. Konermann S, Lotfy P, Brideau NJ, Oki J, Shokhirev MN, Hsu PD. Transcriptome Engineering with RNA-Targeting Type VI-D CRISPR Effectors. Cell. 2018;173(3):665–676 e614.

44. Morelli KH, Wu Q, Gosztyla ML, et al. An RNA-targeting CRISPR-Cas13d system alleviates disease-related phenotypes in Huntington’s disease models. Nat Neurosci. 2023;26(1):27–38.

45. Trochet D, Prudhon B, Vassilopoulos S, Bitoun M. Therapy for dominant inherited diseases by allele-specific RNA interference: successes and pitfalls. Curr Gene Ther. 2015;15(5):503–510.

46. Battistella G, Simonyan K. Clinical Implications of Dystonia as a Neural Network Disorder. Adv Neurobiol. 2023;31:223–240.

47. Taiwo FT, Adebayo PB. Neuroimaging findings in DYT1 dystonia and the pathophysiological implication: A systematic review. Brain Behav. 2023;13(6):e3023.

48. Wu J, Ren J, Luo H, Zuo X, Tang Y. Generation of patient-specific induced pluripotent stem cell line (CSUi002-A) from a patient with isolated dystonia carrying TOR1A mutation. Stem Cell Res. 2021;53:102277.

49. Tang Y, Ren J, Li CC. Establishment of a GFP::LMNB1 knockin cell line (CSUi002-A-1) from a dystonia patient-specific iPSC by CRISPR/Cas9 editing. Stem Cell Res. 2021;55:102505.

50. Akter M, Cui H, Chen YH, Ding B. Generation of gene-corrected isogenic control cell lines from a DYT1 dystonia patient iPSC line carrying a heterozygous GAG mutation in TOR1A gene. Stem Cell Res. 2022;62:102807.

51. Akter M, Cui H, Chen YH, Ding B. Generation of two induced pluripotent stem cell lines with heterozygous and homozygous GAG deletion in TOR1A gene from a healthy hiPSC line. Stem Cell Res. 2021;56:102536.

