## Supplementary material for "Allele-specific targeting of mutant TOR1A by the compact CRISPR/NmCas9 system in DYT1 dystonia with high fidelity": Supplementary Figures.docx

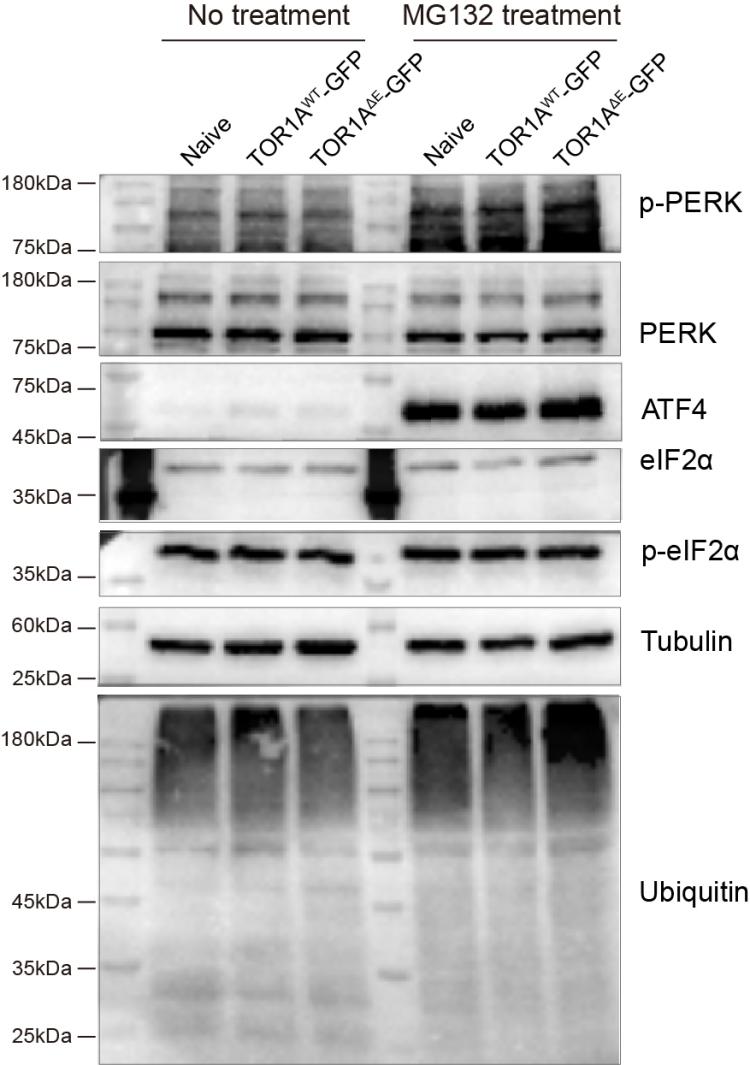

**Supplementary Figure 1 The protein quality control machinery and ER stress in knockin cells.** To examine the effects of the mutant TOR1A in generated knockin cells, both heterogeneous TOR1A-2A-GFP cells with treated with MG132. The expression of PERK-eIF2α signaling genes associated with ER stress and the accumulation of ubiquitin were detected by western blot.

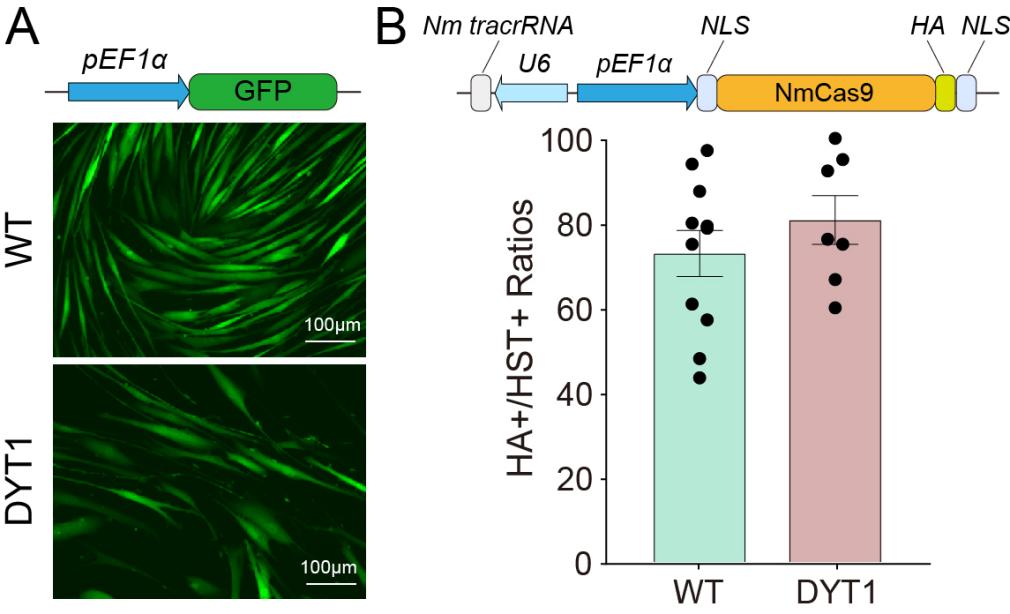

**Supplementary Figure 2 Transduction efficiency of lentiviruses in fibroblasts.** (A) Transduction with the control lentivirus (pEF1α-GFP). Scale bars, 100 μm. (B) Transduction efficiencies of NmCas9/sgMut-Nm21/24 lentiviruses were assessed by immunostaining with HA.
